# The neurobiology of BRD1 implicates sex-biased dysregulation of nuclear receptor signaling in mental disorders

**DOI:** 10.1101/257170

**Authors:** Anto P. Rajkumar, Per Qvist, Sanne H. Larsen, Ross Lazarus, Jonatan Pallesen, Nicoletta Nava, Gudrun Winther, Nico Liebenberg, Veerle Paternoster, Tue Fryland, Johan Palmfeldt, Kim Fejgin, Arne Mørk, Mette Nyegaard, Bente Pakkenberg, Michael Didriksen, Jens R. Nyengaard, Gregers Wegener, Ole Mors, Jane H. Christensen, Anders D. Børglum

**Affiliations:** iPSYCH, The Lundbeck Foundation Initiative for Integrative Psychiatric Research, Denmark; Department of Biomedicine, Aarhus University, Aarhus, Denmark; Centre for Integrative Sequencing, iSEQ, Aarhus University, Aarhus, Denmark; Mental Health of Older Adults and Dementia Clinical Academic Group, South London and Maudsley NHS foundation Trust, London, UK; Department of old age psychiatry, Institute of Psychiatry, Psychology, & Neuroscience, King’s College London, London, UK; Synaptic transmission, H. Lundbeck A/S, Copenhagen, Denmark; Computational Biology, Baker IDI heart and diabetes institute, Melbourne, Australia; Research Unit for Molecular Medicine, Aarhus University Hospital, Denmark; Translational Neuropsychiatry Unit, Department of Clinical Medicine, Aarhus University Hospital, Aarhus, Denmark; Research Laboratory for Stereology and Neuroscience, Bispebjerg University Hospital, Copenhagen, Denmark; Core Centre for Molecular Morphology, Section for Stereology and Microscopy, Department of Clinical Medicine, Centre for Stochastic Geometry and Advanced Bioimaging, Department of Clinical Medicine, Aarhus University, Denmark; Psychosis Research Unit, Department of Clinical Medicine, Aarhus University Hospital, Aarhus, Denmark.

## Abstract

The schizophrenia and bipolar disorder associated gene, *BRD1*, encodes a scaffold protein that in complex with epigenetic modifiers regulate gene sets enriched for psychiatric disorder risk. Preclinical evidence from male *Brd1*^+/−^ mice has previously implicated BRD1 with phenotypes of translational relevance to schizophrenia. Here we describe the phenotype of female *Brd1*^+/−^ mice and report attenuated dendritic architecture and monoaminergic dysregulation accompanied by sex-specific changes in affective behaviors. In accordance, global gene expression profiling reveals regional dysregulation of gene sets enriched with major depressive disorder and schizophrenia risk in female and male *Brd1*^+/−^ mice, respectively. Independent of sex, however, differentially expressed genes cluster in common functional pathways associated with psychiatric disorders, including mitochondrial dysfunction and oxidative phosphorylation as well as G-protein coupled-, and nuclear receptor mediated signaling. Accordingly, we provide *in vitro* evidence that BRD1 modulates the transcriptional drive of a subset of nuclear receptors (e.g. the vitamin D and glucocorticoid receptors). Moreover, we demonstrate enrichment of psychiatric disorder risk in the target genes of nuclear receptors, sex-biased expression of several nuclear receptor genes in the adult brain of *Brd1*^+/−^ mice, and that sex-biased genes in general are enriched with nuclear receptor genes particularly at the earliest developmental stage of the human brain. Overall, our data suggests that the spatio-temporal interaction between BRD1 and subsets of nuclear receptors in the brain is sex-biased and that hampered BRD1 mediated regulation of target genes governed by certain nuclear receptors may significantly contribute to sex differences in psychopathology.

## Introduction

Psychiatric disorders are complex multifactorial illnesses characterized by shared genetic risk ^1^, overlap in clinical profiles and documented sex differences in their prevalence, symptomatology, and course ^2–6^. Epigenetic processes, such as acetylation of histone lysine residues, are linked with brain development as well as lifelong neural plasticity ^7^ and have been implicated with the pathophysiology of both psychotic and affective disorders ^8^. Bromodomain containing-1 (BRD1) has been identified in complexes involved with histone acetylation and chromatin remodeling ^9,10^ and interacts at genomic sites enriched with genes implicated in neurodevelopmental processes ^10^. *BRD1* is widely expressed in human brain ^11^, differentially regulated in limbic and neocortical tissues upon exposure to external stressors in rats ^12,13^, and involved in the epigenetic regulation of embryonic development, survival, and differentiation of embryonic stem cells ^9,14,15^. Supporting a role for *BRD1* in mental health, *BRD1* has repeatedly been associated with schizophrenia and bipolar disorder in large genetic studies ^16–20^, including gene-wise significant association in the currently largest schizophrenia GWAS mega-analysis ^21^ and genome-wide significance in the Psychiatric Genomics Consortium (PGC1) schizophrenia sample using an Empirical Bayes statistical approach ^22^. Furthermore, a schizophrenia case with a disruptive nonsense mutation in *BRD1,* which is generally highly intolerant to loss of function mutations ^23^, has been reported ^24^. We have recently shown that male mice with reduced expression of *Brd1* (*Brd1*^+/–^ mice) recapitulate features relating to schizophrenia symptomatology ^25,26^ (**Table 1**). In the present study we assess the impact of reduced *Brd1* expression on sex differences in the behavioral, neurochemical, and neurostructural domains in mice. Through global gene expression profiling of selected brain structures and integrative analyses of human datasets, we provide evidence linking functional pathways and molecular mechanisms to sex differences in neuropsychiatric pathology.

**Table 1.** Basic neurological functioning and behaviors in *Brd1*^+/−^ mice.

| Test | Parameters | ♂ | ♀ | Implication: |
| --- | --- | --- | --- | --- |
| **Irwin's observational battery** | Undisturbed behavior | - | - | Basic neurological functioning |
|  | Finger approach | - | - | Basic neurological functioning |
|  | Touch escape | - | - | Basic neurological functioning |
|  | Grip strength | - | ↓ | Basic neurological functioning |
|  | Visual placing response | - | - | Basic neurological functioning |
|  | Corneal response | - | - | Basic neurological functioning |
|  | Toe-pinch response | - | - | Basic neurological functioning |
|  | Wire-maneuver | - | ↓ | Basic neurological functioning |
|  | Limb- and abdominal tone | - | - | Basic neurological functioning |
|  | Tail-pinch response | - | - | Basic neurological functioning |
| **Hot-plate** | Response | - | - | Acute pain response |
| **Hidden food retrieval** | Time to retrieve | - | ND | Olfactory functioning |
| **Beam walking** | Crossing speed/missteps | - | - | Motor coordination |
| **Rota-rod** | Latency to fall | - | ↓ | Motor coordination |
| **Foot-printing test** | Stride length | - | - | Motor coordination |
|  | Base width | - | - | Motor coordination |
|  | Step uniformity | - | ↓ | Motor coordination |
| **Social interaction** | Passive interaction | ↓ | ND | Associability |
|  | Aggressive interaction | ↑ | ND | Aggression |
|  | Latency to interaction | ↑ | ND | Social withdrawal |
| **3 chamber test** | Sociability | ↓ | ND | Social withdrawal |
|  | Social recognition | - | ND | Social cognition |
|  | Remote social memory | ↓ | ND | Long term recognition memory |
| **Spontaneous alternation (SA)** | Baseline | - | ND | Working memory |
|  | PCP induced | ↓ | ND | Working memory |
| **Continuous alternation (CA)** | Baseline | - | ND | Working memory |
|  | PCP induced | ↓ | ND | Working memory |
| **Fear Conditioning (FCS)** | Conditioning | ↓ | ↓ | Conditional learning |
|  | Contextual memory (day 2) | ↓ | ↓ | Associative memory |
|  | Contextual memory (day 3) | ↓ | ND | Associative memory |
|  | Contextual memory (day 7) | ↓ | ND | Associative memory |
|  | Extinction retrieval | - | - | Associative memory |
|  | Cue dependent learning | ↓ | - | Associative memory |
| **Acoustic startle reactivity (ASR)** | Startle | ↑ | ↑ | Hearing/stress susceptibility |
|  | Latency to startle | ↓ | ↓ | Stress susceptibility |
| **Prepulse inhibition (PPI)** | Baseline | - | ↓ | Pre-attentive processing |
|  | PCP induced | ↓ | ND | Pre-attentive processing |
|  | Amphetamine induced | - | ND | Pre-attentive processing |
| **Locomotor activity** | Novelty-induced | - | - | Psycho-motor activity |
|  | PCP induced | ↑ | ND | Cortico-thalamic/Meso-limbic drug responsiveness |
|  | Amphetamine induced | - | - | Meso-limbic drug responsiveness |
|  | Cocaine-induced | ↑ | - | Meso-limbic drug responsiveness |
| **Delayed alternation (DA)** |  | ↓ | ND | Working memory/spatial reference memory |
| **8 arm radial maze (ARM)** | Re-entry to baited arms | ↑ | - | Working memory |
|  | Entry to non-baited arms | ↑ | ↑ | Non-spatial reference memory |
| **Morris water maze (MWM)** | Acquisition | - | ND | Learning |
|  | Probe test | - | ND | Spatial reference memory |
|  | Flag test | - | ND | Vision |
| **Attentional set shifting (ASST)** | Rule learning | ↓ | ND | Learning |
|  | Reverse learning | - | ND | Reverse learning |
|  | Intradimensional shift | - | ND | Executive functioning |
|  | Extradimensional shift | ↓ | ND | Executive functioning |
| **Elevated plus maze (EPM)** | Time in open arms | - | - | Anxiety behavior/Mania |
| **Bright open field (BOF)** | Time in central zone | - | - | Anxiety behavior/Mania |
| **Light and dark box (LDB)** | Time in light box | ↓ | - | Anxiety behavior/Mania |
| **Open field test (OF)** | Distance moved | - | - | Anxiety behavior/Mania |
| **Forced swim test (FST)** | Immobility | - | ↑ | Behavioral despair/Mania |
| **Tail suspension test (TST)** | Immobility | - | ↑ | Behavioral despair/Mania |
| **Sucrose preference test (SPT)** | Sucrose preference | ND | ↓ | Anhedonia |
| **PTZ induced seizure activity** | Myoclonic jerks (#) | ↑ | ND | Sensitivity of GABA A receptor response (local) |
|  | Clonic seizures (#) | ↑ | ND | Sensitivity of GABA A receptor response (global) |
|  | Clonic seizures (onset) | ↓ | ND | Sensitivity of GABA A receptor response (global) |
|  | Clonic-tonic seizures | - | ND | Sensitivity of GABA A receptor response (global) |

Data presented in this manuscript are highlighted in grey, whereas other data have previously been reported on <sup>25,26</sup>. ND: not determined

## Materials and Methods

### Animals

A mouse line heterozygous for a targeted deletion within the *Brd1* gene, *Brd1*^tm1569_4.2Arte^ (*Brd1*^+/−^) was generated by TaconicArtemis GmbH (Cologne, Germany) using a targeting vector (pBrd1 Final cl 1 (UP0257)) with loxP sites flanking exon 3 to 5 of the *Brd1* gene. For further details, see **Supplemental Methods and Materials**.

### General Assessment of Neurology, Motor Coordination and Behavioral tests

Details on functional observation battery, acute pain response, rotarod, balance beam walking, footprinting, forced swim test (FST), tail suspension test (TST), sucrose preference test (SPT), open field (OF), bright open field (BOF), light and dark box (LDB), elevated plus maze (EPM), fear conditioning (FC), 8-arm radial maze (8ARM), 24 hours locomotion (24HLM), Prepulse inhibition (PPI) and amphetamine (AIH) or cocaine induced hyperactivity (CIH) can all be found in **Supplemental Methods and Materials**.

### Experimental design

All experiments involved 7-15 mice, 8-11 weeks old, in each group (figure legends present exact numbers). Observer was blind to mouse genotypes. One batch of mice completed OF, TST and FST, while another completed BOF, LDB and EPM. Mice were not reused for other experiments. In this study, general assessment of neurology and testing in motor coordination tests, PPI, FC, 8ARM, AIH, CIH, SPT, and 24HLM were only performed in female mice. Other tests were completed by both male and female mice. All studies were carried out in accordance with Danish legislation, and permission for the experiments was granted by the animal welfare committee, appointed by the Danish Ministry of Food, Agriculture and Fisheries – Danish Veterinary and Food Administration.

### Quantification of neurotransmitters

Mice were sacrificed by cervical dislocation and frontal cortical-, hippocampal-, and striatal tissues were collected by free-hand dissection and processed for quantitative high-pressure liquid chromatography (HPLC) analyses of dopamine and serotonin. For details on HPLC procedures, see **Supplemental Methods and Materials**.

### Brain morphometry, Golgi-cox staining and 3-D image analysis

Left cerebral hemispheres (n=8/group) were stained with FD Rapid Golgi-Stain kit (FD Neurotechnologies, Ellicott City, USA), and cut into 150 µm thick-slices on a vibratome-3000 (Vibratome, St Louis, MO, USA). Anterior cingulate cortex (aCC) pyramidal neurons were identified (60X; oil-immersion; numerical-aperture=1.4) by their prominent apical dendrites, and 6 neurons/mouse were chosen by the optical disector. Image stacks (90-105 consecutive images at 1 µm interval) were captured by optical wide-field microscopy (Olympus BX50, Tokyo, Japan) and newCAST software (Visiopharm, Hoersholm, Denmark). 3-D image reconstruction and analyses were completed using Imaris software version 7.6.3 (Bitplane AG, Zurich, Switzerland). For description on brain morphometric analyses, see **Supplemental Methods and Materials**.

### RNA-sequencing and data analyses

Mouse brains (n=8/group) were sectioned coronally (1 mm thick) using a slicer matrix (Zivic Instruments, Pittsburgh, USA). Right amygdala, striatum, aCC and hippocampus CA3 were identified, and punched by a punch-needle (1 mm diameter) at −20°C. RNA was extracted using Maxwell-16 instrument system and LEV simplyRNA Tissue Kit (Promega, Madison, USA). cDNA synthesis with random hexamer primers and TruSeq library preparation, RNA-sequencing (50bp; single-end; minimum 10 clean million reads/sample) was performed using Illumina HiSeq2000 (Illumina, San Diego, USA). Reads were aligned to mouse genome (Mus_musculus.GRCm.38.72) by TopHat2.0.6 and counted by HTSeq0.5.4. Differentially Expressed Genes (DEGs) were identified by edgeR3.2.4. For complete description of RNA-sequencing and data analyses, see **Supplemental Methods and Materials**.

### Generation of BRD1^CRISPRex6/+^ HEK cell lines and Nuclear Receptor 10-Pathway Reporter Array

The CRISPR/Cas9 system was used to establish *BRD1* knock down human embryonic kidney (HEK) cell lines (HEK *BRD1*^CRISPRex6/+^) and their transcriptional drive was investigated in a Cignal Finder Nuclear Receptor 10-Pathway Reporter Arrays (Qiagen). Briefly, 4 different colonies of *BRD1*^CRISPRex6/+^ and 4 colonies of naïve HEK cells were co-transfected with plamids carrying a firefly luciferase reporter coupled to individual NR transcriptional response elements (12.5 ng / 1000 cells) and a plasmid carrying a renilla reporter coupled to a constitutively active cytomegalovirus (CMV) promoter (12.5 ng / 1000 cells) using 0.2 μL TurboFect^TM^ Transfection Reagent (ThermoScientific). Cells were then cultured for 48 hours followed by cell lysis. Firefly and renilla luciferase activity was measured using the Dual-Luciferase® Reporter (DLRTM) Assay System (Promega) on a MicroLumat Plus LB96V (Berthold Technologies, Bad Wildbad, Germany) according to manufacturer’s protocol. For further details, see **Supplementary Methods and Materials**.

### Analysis of disease risk enrichment in DEG sets and in nuclear receptor response element containing genes

Gene set analysis was performed with Multi-marker Analysis of GenoMic Annotation (MAGMA) ^27^ using default settings, based on summary statistics from selected publicly available GWASs while excluding the MHC region and imputed SNPs with info score < 0.8. For the analysis of NR response element containing genes, available human MEME format motifs corresponding to curated, non-redundant sets of profiles for NR monomers and heterodimers were downloaded from The JASPAR CORE database (jaspar.genereg.net/) and genomic positions were identified using the FIMO tool (http://meme-suite.org/tools/fimo). Gene lists were generated for genes containing their respective DNA motifs within a broadly defined promoter region spanning transcription start site (TSS) and 2000 bp upstream. See **Supplemental Methods and Materials** for details.

## Results

### General assessment of neurology and motor coordination

Male and female *Brd1*^+/−^ mice were overall healthy, as described elsewhere ^25^. However, female mice showed marginally impaired growth, slightly reduced size, and a near significant reduction in blood glucose concentration ^25^. Systematic testing of general neurological functions in female *Brd1*^+/−^ mice, revealed slightly reduced performance in the grip strength and wire-maneuvering tasks (**Table 1**). Female *Brd1*^+/−^ mice showed normal pain response (**Figure S1A**) but were mildly impaired in their motor coordination as evident from their rotarod performance (**Figure S1B**, *p*=0.047)) and gaiting pattern (**Figure S1F**, gaiting uniformity, *p*=0.039). However, as they performed *at par* with their WT littermates in the beam walking task (**Figure S1G-H**), we considered female *Brd1*^+/−^ mice fit for testing in settings assessing complex behaviors. Assessment of motor coordination in male *Brd1*^+/−^ mice has been reported on previously ^25^.

### Assessment of affective behaviors

General locomotor activity was assessed in the OF where male (**Figure 1A**) and female (**Figure 1B**) *Brd1*^+/−^ mice performed *at par* with their WT littermates. Risk taking behaviors were assessed in the BOF, LDB and EPM revealing no consistent differences in performance between *Brd1*^+/−^ and WT mice (see **Supplementary Results** for further details). Circadian rhythm, measured as their 24HLM performance, appeared unaltered in female *Brd1*^+/−^ mice (**Figure 1C**). Similar to what has previously been reported in male *Brd1*^+/−^ mice ^25^, female *Brd1*^+/−^ mice displayed significantly increased acoustic startle responsivity (ASR) (**Figure 1D**, *p*=0.003), both when initially introduced to the test setting (**Figure 1D**, *p*=0.004) and before baseline PPI testing (**Figure 1D**, *p*=0.032). Response latency to the startle was furthermore significantly shorter in female *Brd1*^+/−^ mice than in WT mice (**Figure 1E**, *p*=0.044). Interestingly, female *Brd1*^+/−^ mice displayed reduced PPI across the span of tested prepulse intensities (**Figure 1F**, *p*=0.049), whereas this has only been reported in male *Brd1*^+/−^ mice at a high prepulse intensity and after administration of the psychostimulatory drug, phencyclidine (PCP) ^25^. Suggestive of behavioral despair, female *Brd1*^+/−^ mice were significantly more immobile in TST (**Figure 1G,** *p*=0.007) and in FST (**Figure 1H**, *p*=0.002) compared to WT mice, whereas this was not evident in male *Brd*1^+/−^ mice (**Figure 1I** and **Figure 1J**). Sucrose preference was similarly significantly reduced in female *Brd1*^+/–^ mice compared to WT mice (**Figure 1K,** *p*=0.001).

**Figure 1:**
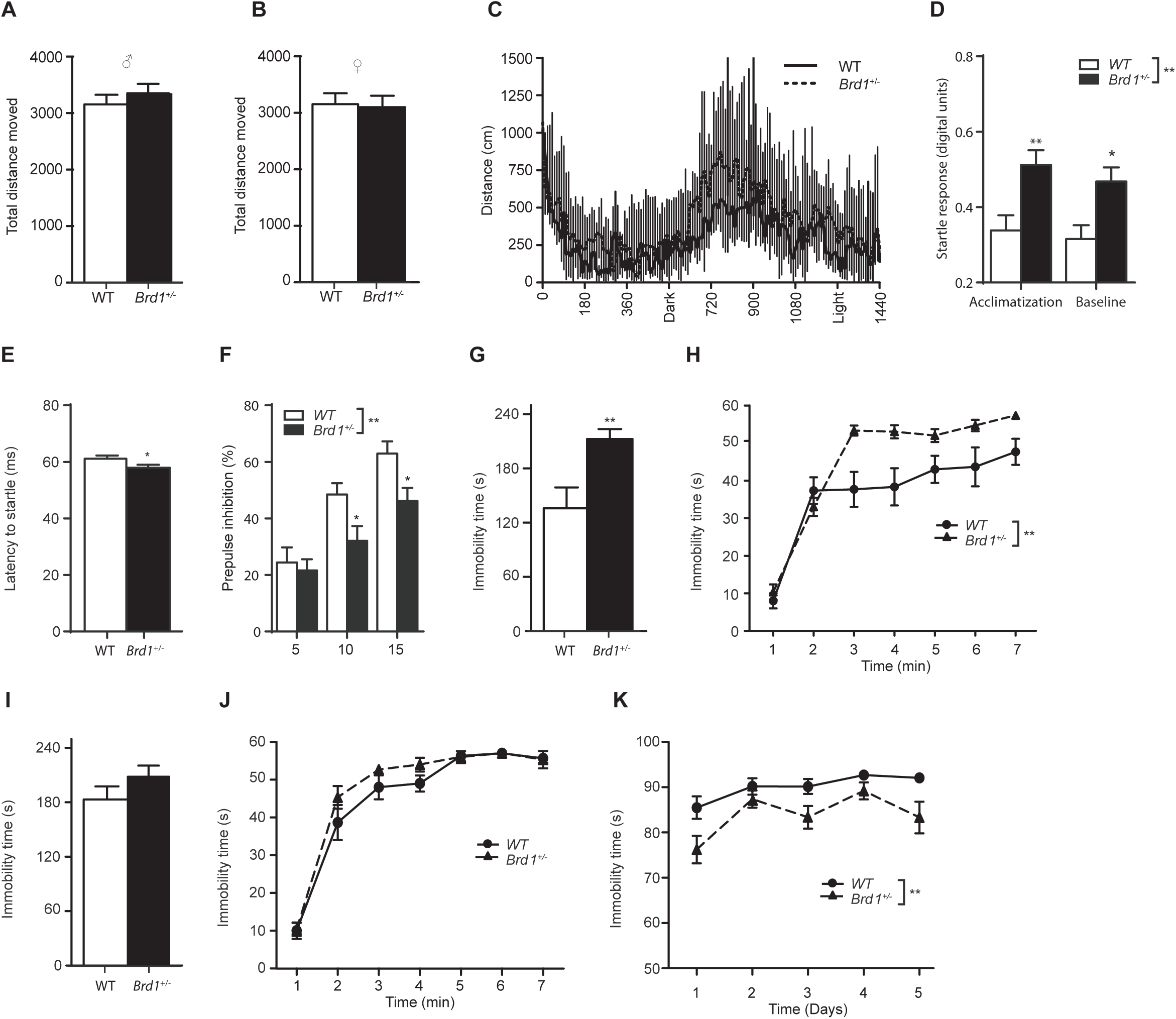
Behavioral characterization in male and female *Brd1*^+/−^ mice. A) Male mice (Mann-Whitney U=94.0; *p*=0.46); **B)** female mice (t=0.20; *p*=0.85): Total distance moved in the Open Field (OF) (n=15/group); **C)** Female mice: Distance moved over 24 hours (n=10/group, F=1.11; *p*=0.31). Dark indicates the time, when the lights were switched off in the stable, while light indicates the time, when they were switched on; **D)** Female *Brd1*^+/−^ mice (n=18) displayed significantly increased acoustic startle responsivity (ASR) compared to WT mice (n=17) (genotype effect, F=10.10, *p*=0.003) both when initially introduced to the test setting (Tukey’s *post hoc* test, *p*=0.004) and before baseline PPI testing (Tukey’s *post hoc* test, *p*=0.032); **E)** Response latency to the startle was furthermore significantly shorter than in WT mice (t= 2.09, *p*=0.044); **F)** Female *Brd1*^+/−^ mice (n=18) displayed reduced prepulse inhibition (PPI) compared to WT mice (n=17) across the span of tested prepulse intensities (genotype effect, F=4,163, *p*=0.049); **G)** Female *Brd1*^+/−^ mice were significantly more immobile in TST compared to WT mice (n=15/group; t=3.01; *p*=0.007); **H)** Female *Brd1*^+/−^ mice were significantly more immobile in FST compared to WT mice (n=15/group; F=12.26; *p*=0.002); **I)** Male *Brd*1^+/−^ mice performed at par with WT mice in TST (n=15/group); t=1.34; *p*=0.19) and; **J)** in FST (n=15/group; F=3.26; *p*=0.08); **K)** Sucrose preference (weight of 2% sucrose solution consumed/ weight of total fluid consumed) in percentage. Sucrose preference was significantly reduced in female *Brd1*^+/−^ mice compared to WT mice (n=11/group; F=14.03; *p*=0.001). *: p<0.05; **: p<0.01; ***: p<0.001.

### Cognitive behaviors

Like reported in male *Brd1*^+/−^ mice ^25,26^, female *Brd1*^+/−^ mice froze significantly less than WT mice during the conditioning phase of FCS (**Figure S3A,** *p*=0.002) and when returning to the same context on the following day (**Figure S3B,** *p*=0.03), suggesting that female *Brd1*^+/−^ mice have context-dependent learning deficits thereby paralleling the cognitive deficits reported in male *Brd1*^+/−^ mice ^25,26^. However, unlike their male counterparts ^26^, female *Brd1*^+/−^ mice did not display cue dependent memory deficits in FCS (**Figure S3C**) and did not differ in working memory errors in 8ARM (**Figure S3D**). However, they made significantly more entries into the never-baited arms (**Figure S3E,** *p*=0.03), suggesting impaired reference memory, like also reported in male *Brd1*^+/−^ mice ^26^.

### Neurochemistry and psychotropic drug-induced activity

As has previously been reported in male *Brd1*^+/−^ mice ^25^, female mice displayed unaltered hippocampal serotonin levels (**Figure 2A, B**), significantly reduced hippocampal dopamine levels (**Figure 2A, C**, *p*=0.045) and unaltered fronto-cortical dopamine levels (**Figure 2A, D**). However, contrary to male *Brd1*^+/−^ mice ^25^, female mice had significantly less fronto-cortical serotonin (**Figure 2A, E,** *p*=0.01) and, noticeably, significantly reduced striatal dopamine (**Figure 2A, F**, *p*=0.02) compared to WT mice. Additionally, their sensitivity towards the psychomotor stimulatory effects of amphetamine 5 mg/kg (**Figure 2G**) and cocaine 15 or 30 mg/kg (**Figure 2H**) did not differ from the sensitivity of WT mice.

**Figure 2:**
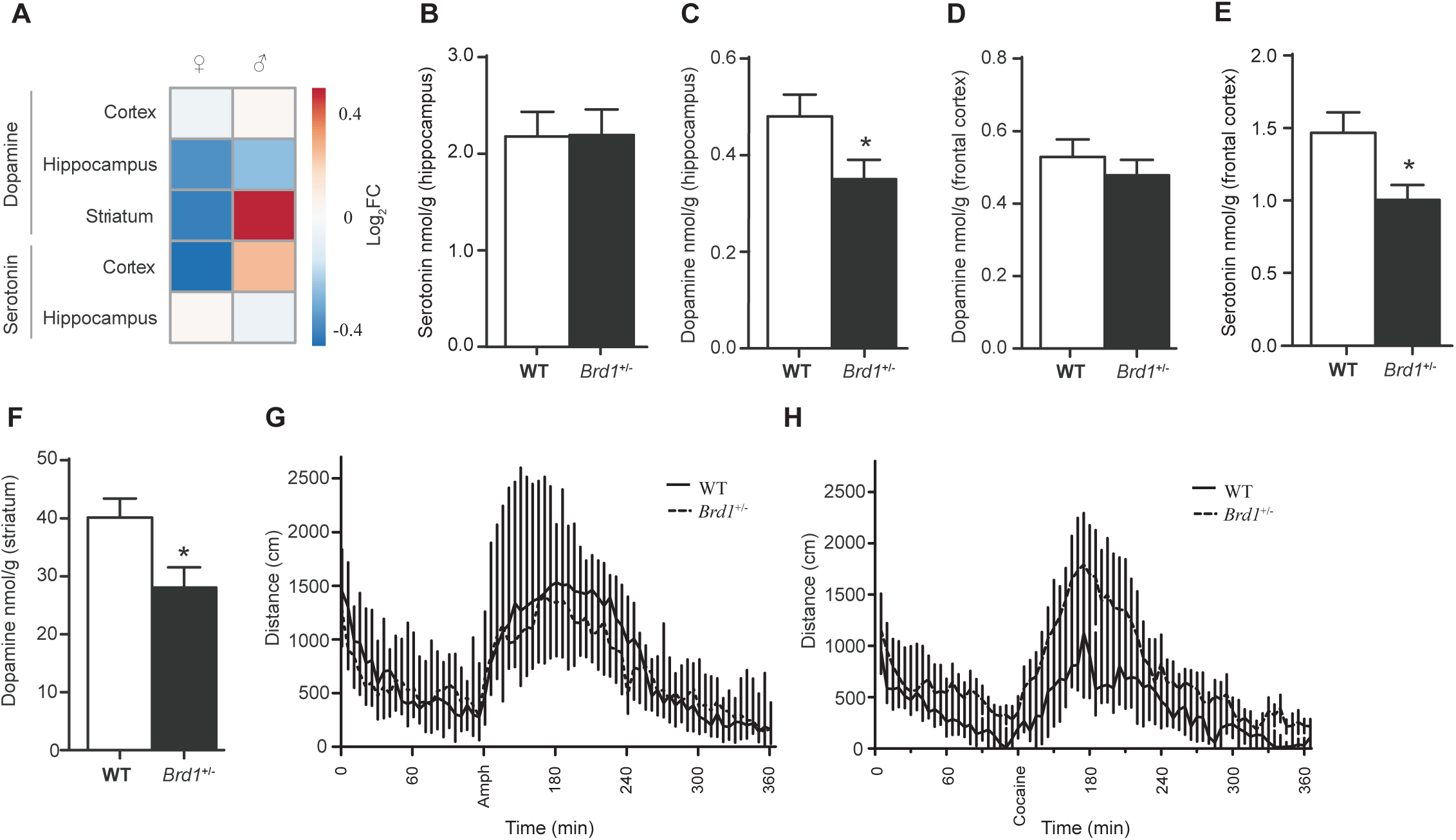
Neurochemistry and psychotomimetic drug sensitivity in female *Brd1*^+/−^ mice. **A-E)** Neurotransmitter levels were determined by HPLC in several brain tissues in *Brd1*^+/−^ mice**; A)** Front: Log2 fold changes (Log_2_FC) in monoamine levels between WT and *Brd1*^+/−^ female mice aligned with corresponding data for male mice as previously reported ^25^; **B)** Female mice displayed unaltered hippocampal serotonin level (n=9/group; t=0.043, *p*=0.966); **C)** significantly reduced hippocampal dopamine level (n=10/group, t=2.147, *p*=0.045); **D)** unaltered fronto-cortical dopamine (n=9/group; t=0.788, *p*=0.441); **E)** less fronto-cortical serotonin (n=15/group; t=2.70; *p*=0.01) and; **F)** reduced striatal dopamine (n=10/group; t=2.52; *p*=0.02) compared to WT mice. 5-HT: 5-Hydroxy Tryptamine (Serotonin); DA: Dopamine; **G)** Distance moved before and after amphetamine 5 mg/kg (Amph) injection was similar in female *Brd1*^+/−^ and WT mice (n=10/group; F=1.12; *p*=0.30); **H)** Distance moved before and after cocaine 30 mg/kg injection was similar in female *Brd1*^+/−^ and WT mice (n=12/group; F=1.91; *p*=0.25). *: p<0.05

### Brain volume and neuronal morphology

Total brain volume, as estimated by stereology, was slightly reduced (~8%) in female *Brd1*^+/−^ mice (**Figure 3A** and **Figure S4A,** *p*=0.041), but with no difference in brain symmetry (**Figure S4B**) or ventricle volume (**Figure S4C-D**). In line with reduced overall brain tissue volume, aCC pyramidal neurons had significantly shorter dendrites in female *Brd1*^+/−^ mice compared to WT mice (**Figure 3B,** *p*=0.008) combined with less dendritic branching (**Figure 3C,** *p*=0.01) and less dendritic spine density (**Figure 3D,** *p*<0.001). 3-D Sholl analysis counting the dendritic intersections on the concentric spheres with their centres at soma confirmed that these neurons had significantly less dendritic branching (**Figure 3E-F,** *p*<0.001).

**Figure 3:**
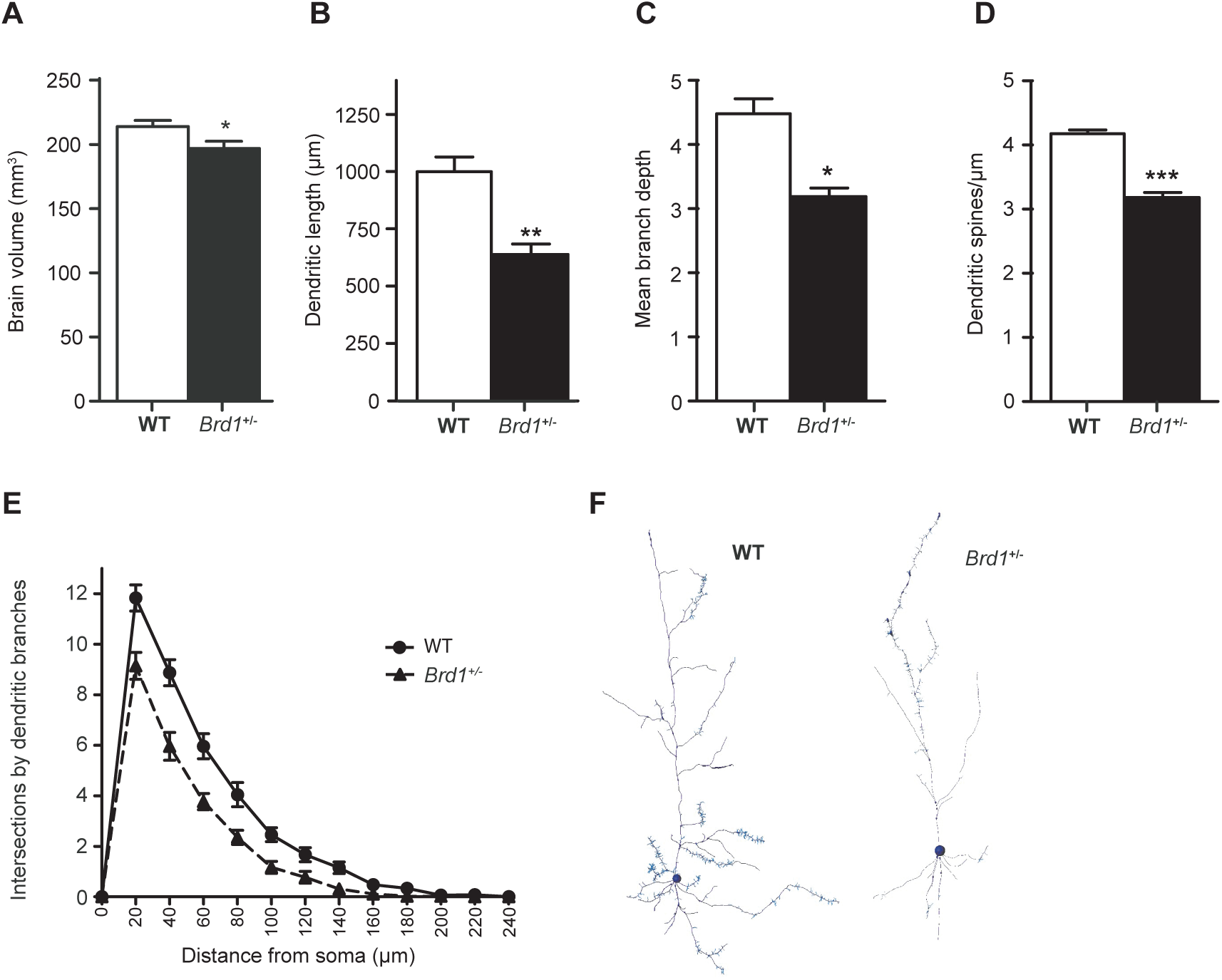
Brain morphometry in female *Brd1*^+/−^ mice. **A)** Total brain volume was slightly reduced in female *Brd1*^+/−^ mice compared to WT mice (n=7/group; t=2.31; *p*=0.041); **B)** Total dendritic length including apical and basal dendrites. aCC pyramidal neurons had significantly shorter dendrites in female *Brd1*^+/−^ mice compared to WT mice (t=3.29; *p*=0.008); **C)** Mean branch depth: Branching depth was defined by the number of bifurcations from the beginning point to the end of a dendrite. Female *Brd1*^+/−^ mice had less dendritic branching (t=3.08; *p*=0.01) compared to WT mice; **D)** Mean dendritic spine density (number of spines/ length of dendrites). Female *Brd1*^+/−^ mice had less dendritic spine density (t=9.19; *p*<0.001) compared to WT mice; **E)** 3-D Sholl analysis: Number of dendritic intersections on concentric spheres (radius interval 20µm) with their centres at soma. Neurons in female *Brd1*^+/−^ mice had significantly less dendritic branching (F=20.60; *p*<0.001) than neurons in WT mice; **F)** 3-D reconstruction of left: WT neuron and right: *Brd1*^+/−^ neuron *: p<0.05; **: p<0.01; ***: p<0.001.

### Global gene expression profiling, pathway analyses and assessment of disease risk enrichment

To delineate the molecular signatures accompanying the behavioral-, neurochemical-, and brain morphometric changes identified in female *Brd1*^+/−^ mice, we conducted global gene expression profiling of selected brain tissue micropunches. All tissues were characterized by high numbers of nominally significant DEGs (**Table S1-5**). However, whereas 82 DEGs were significant after Benjamini-Hochberg false discovery rate (FDR) correction at 5% in amygdala, only three DEGs were identified in the aCC, including *Brd1*, which again was the only DEG detected in hippocampus (CA3) and striatum. Hence, downstream analyses were conducted on nominally significant DEGs (for data on validation of DEGs, see ^28^). Despite the limited overlap of DEGs between the regions (**Figure 4A**), nominally significant DEGs clustered in partly overlapping functional pathways (**Figure 4B-E**) converging in G-protein coupled receptor (GPCR)-cAMP-DARPP-32 signaling in cortex, hippocampus, and amygdala (**Figure 4B, C and E**). Immuno-signaling pathways were affected in hippocampus, striatum, and amygdala (**Figure 4C-E**). Notably, common to all regions was the clustering of DEGs in signaling pathways related to nuclear receptor (NR) mediated signaling, including retinoid acid receptor (RAR)-, vitamin D receptor/retinoid X receptor (VDR/RXR)-, farnesoid X receptor (FXR)/RXR-, liver X receptor (LXR)/RXR, and thyroid hormone receptor (THR) mediated signaling as well as the Hepatic fibrosis /Hepatic stellate cell activation pathway of which VDR, FXR, LXR and retinoids are documented negative modulators ^29^ (**Figure 4B-E**). In line with decreased sucrose preference and despair-like behaviors in female *Brd1*^+/−^ mice, nominally significant cortical DEGs showed significant enrichment for major depressive disorder genetic risk (**Figure 4F,** *p*=0.007), while no risk enrichment was seen for bipolar disorder, schizophrenia (**Figure 4F**) or any of the three assessed non-brain disorders (**Figure S5**).

**Figure 4:**
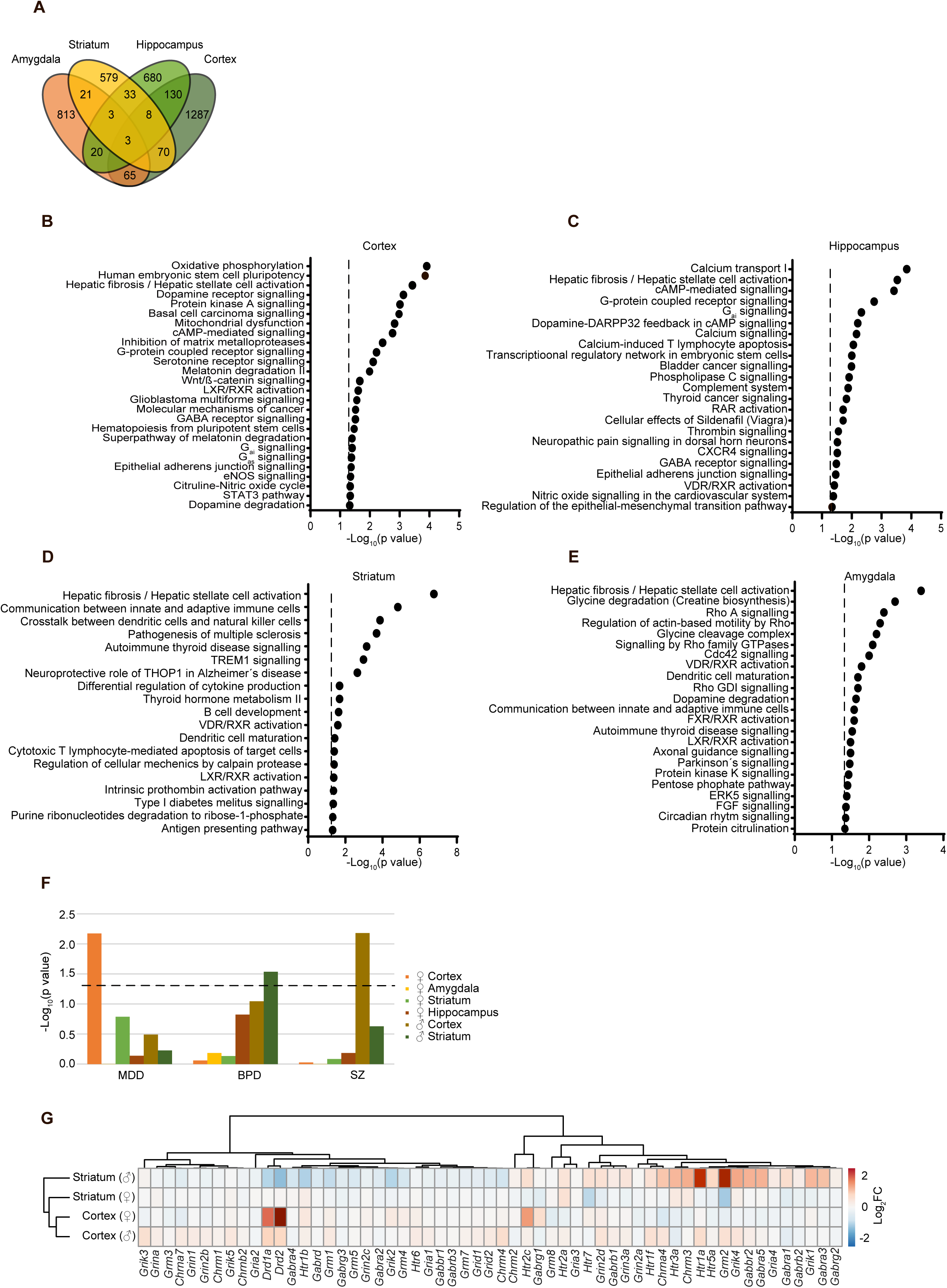
Gene expression profiling and disorder risk enrichment in *Brd1*^+/−^ mice. **A)** Illustration of overlap between nominally significant DEGs identified cortex (aCC), hippocampus (CA3), striatum (CPu) and amygdala in female *Brd1*^+/−^ mice; **B-E**) Ingenuity pathway analyses of nominally significant DEGs in **B)** cortex (aCC); **C)** hippocampus (CA3); **D)** striatum (CPu) and; **E)** amygdala. Dotted line marks nominal significance threshold (*p*<0.05); **F)** Enrichment for disease risk genes in DEG sets. Genetic risk enrichment for major depressive disorder (MDD, 54 770 cases) ^79,93^, bipolar disorder (BPD, 16 731 cases) ^94^ and schizophrenia (SZ, 81 080 cases) ^83^ was investigated in DEG sets. Bars indicate the mean –log_10_(*p* value) of enrichment in DEGs identified in: female cortex; female amygdala; female striatum; female hippocampus; male cortex; and male striatum with color codes as indicated. Dotted line marks nominal significance threshold (*p*<0.05) **G)** Heatmap showing log fold change in expression (*Brd1*^+/−^ vs WT mice) for both male and female mice in various brain tissues.

### Comparison of expression profiles between male and female Brd1^+/−^ mice

Utilizing previously published RNA sequencing data from male *Brd1*^+/−^ cortex and striatum ^25^ we compared female and male nominally significant DEGs called using the same DEG analysis pipeline (**Table S6** and **Table S7**). The number of DEGs in male *Brd1*^+/−^ mice was higher than in female mice (striatum: 2815 vs 1577, and cortex: 2679 vs 725, respectively) (**Figure S6**). Nominally significant cortical DEGs overlapped significantly in female and male *Brd1*^+/−^ mice (**Figure S6A**, *p*=0.0366), whereas this was not the case for striatal DEGs (**Figure S6B**). In agreement with the observed regional sex differences in neurochemistry in *Brd1*^+/−^ mice mentioned above, fold changes in expression of genes encoding receptors for neurotransmitters, and particularly receptors for monoamines, were markedly different between female and male *Brd1*^+/−^ mice in both cortical and striatal tissue (**Figure 4G**). Furthermore, contrary to cortical DEGs in female mice, cortical DEGs were significantly enriched with schizophrenia and bipolar disorder risk genes in male mice (**Figure 4F,** *p*=0.009). However, among cortical DEGs identified in both male and female *Brd1*^+/−^ mice, numerous genes are implicated in neural signaling, including significantly increased expression of *Adcy5,* encoding a member of the membrane-bound adenylyl cyclase enzymes responsible for the synthesis of cAMP; *Tacr3,* encoding the receptor for the tachykinin neurokinin 3; *Cacng4,* encoding a subunit of a Calcium Voltage-Gated Channel, as well as reduced expression of the genes encoding the GABA producing enzyme, *Gad2* and the calcium buffer, parvalbumin (*Pvalb*) expressed by a subtype of GABAergic interneurons. Furthermore, as observed in the female *Brd1*^+/−^ mice, cortical DEGs in male mice clustered in functional pathways relating to oxidative phosphorylation and mitochondrial dysfunction (**Figure S7A-B)**. Complementing the clustering of female DEGs in NR related signaling pathways, male cortical and striatal DEG sets both cluster in estrogen receptor (ESR) mediated signaling pathways and biosynthesis pathways (Mevalonate pathway 1 and cholesterol biosynthesis pathway through lanosterol) of which intermediates serve as activating ligands for LXR ^30^ and FXR ^31,32^. (**Figure S7A-B**).

### Nuclear receptor mediated signaling in vitro

NRs are a family of ligand-regulated transcription factors that, upon activation by steroid hormones and various other lipid-soluble ligands, regulate gene expression via co-regulator proteins in a tissue, cell and gene specific manner ^33^. BRD1 contains 4 LXXLL signature motifs ^34,35^ found in the majority of NR coactivators ^36^ and additionally a CoRNR box often found in NR co-repressors ^37^ (**Figure 5A**). To investigate whether BRD1 has the potential to modulate the genomic actions of NRs *in vitro,* we firstly generated 4 lines of HEK cells, in which we used the CRISPR/Cas9 system to introduce disruptive mutations in *BRD1* (*BRD1*^CRISPRex6/+^ HEK cells) (**Figure S8**), and then tested in a reporter array if their drive to initiate transcription via a subset of NRs was affected. We found that, in HEK cells with reduced *BRD1* expression, transcription mediated by Hepatocyte nuclear factor (HNF4, *p*=0.046) and Androgen receptor (AR, *p*=0.037) was significantly increased, whereas transcription mediated by Glucocorticoid receptor (GR, *p*=0.042) and VDR (*p*=0.030) was significantly decreased (**Figure 5B**).

**Figure 5:**
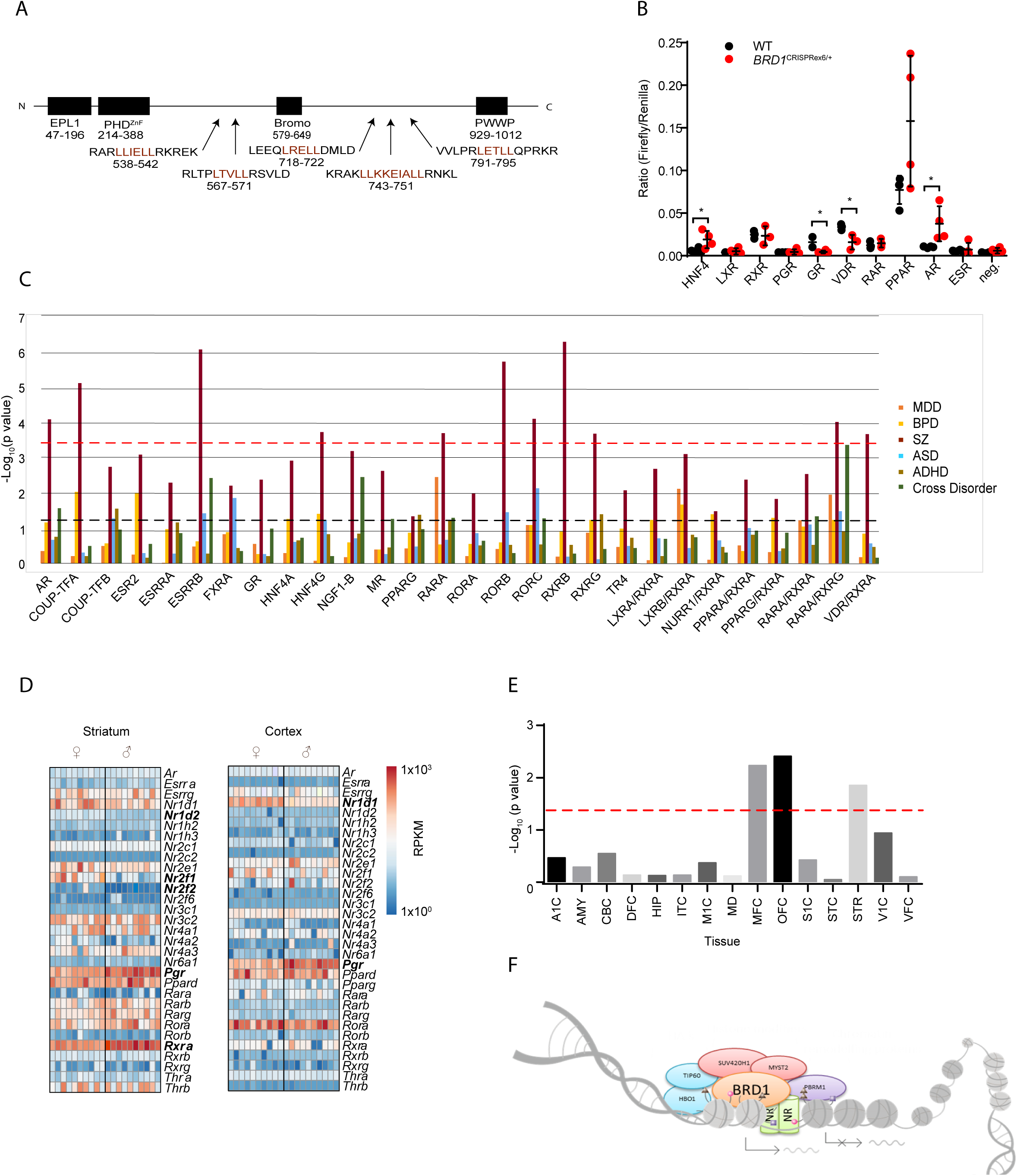
Modulation of nuclear receptor mediated signaling by BRD1, disease risk enrichment in target genes of nuclear receptors and sex-biased expression of nuclear receptor encoding genes in the developing human brain. **A)** Domains and nuclear receptor binding sites in BRD1. Top: Plant homeodomain finger (PHDZnF); Bromodomain (BROMO); Pro-Trp-Trp-Pro (PWWP). Bottom: Amino acid sequences containing putative NR binding sites (4 co-activators (LXXLL) and 1 co-repressor (LXXIXXL)). Pink letters indicate the putative NR binding sites. Amino acid numbering according to BRD1-S ^11^; **B)** Transcriptional drive of 4 distinct *BRD1*^CRISPRex6/+^ clones and 4 WT HEK cell colonies in Dual luciferase array assessing 10 NRs. HNF4 (Hepatocyte nuclear factor); LXR (Liver receptor), RXR (Retinoid X receptor); PGR (Progesterone receptor); GR (Glucocorticoid receptor); VDR (Vitamin D receptor); RAR (Retinoic acid receptor); PPAR (Peroxisome Proliferator Activated Receptor); AR (Androgen receptor); ESR (Estrogen receptor); neg. (negative control (TATA box promoter)). Transcription mediated by Hepatocyte nuclear factor (HNF4; t=2.51; *p*=0.046) and Androgen receptor (AR; t=2.66; *p*=0.037) was significantly increased, whereas transcription mediated by Glucocorticoid receptor (GR; t=2.95; *p*=0.042) and VDR (t=3.31; *p*=0.030) was significantly decreased; **C)** Genetic disease risk enrichment in target genes of various nuclear receptor mono-, and heterodimers. RXRB and G (Retinoid X receptor beta and gamma); RORA, B and C (RAR Related Orphan Receptor alpha, beta and gamma); RARA (Retinoic acid receptor alpha); PPARG (Peroxisome Proliferator Activated Receptor gamma); MR (mineralocorticoid receptor); GR (Glucocorticoid receptor); TR4 (Testicular receptor 4); ESRB (Estrogen receptor beta); ESRRA and B (Estrogen related receptor alpha and beta); AR (Androgen receptor); HNF4A and G (Hepatocyte nuclear factor alpha and gamma); FXRA (Farnesoid X receptor alpha): LXRA (Liver receptor alpha); Coup-TFA and B (Chicken ovalbumin upstream promoter-transcription factor alpha and beta): NURR1 (Nur-related factor 1); LXRA/RXRA; LXRB/RXRA; NURR1)/RXRA; PPARA/RXRA; PPARG/RXRA; RARA/RXRA; RARA/RXRG and VDR (Vitamin D receptor)/RXRA; MDD: Major depressive disorder; BPD: Bipolar disorder; SZ: Schizophrenia; ASD: Autism spectrum disorder; ADHD: Attention deficit hyperactivity disorder; Cross: Cross disorder (MDD, BPD, SZ, ASD and ADHD). The dotted line (dark grey) marks nominal significance threshold (*p*<0.05), red dotted line marks significance threshold after correction for multiple testing (GWASs and NRs); **D)** RPKM values of brain expressed genes encoding nuclear receptors in striatal and cortical tissue in female and male WT mice. Included are 31 genes in striatum and 32 genes in cortex. In striatal tissue, expression of Coup transcription factors (beta, 1 and 2) (*Nr1d2* (t=7.514, *q*<0.001), *Nr2f1* (t=4.551, *q*=0.002), *Nr2f2* (t=3.278, *q*=0.026), the progesterone receptor encoding gene, *Pgr* (t=5.98, *q*=0.032), and *Rxra* (t=5.625, *q*<0.001) differed significantly between female (n=10) and male (n=10) mice after Benjamini-Hochberg false discovery rate (FDR) correction at 5%. In cortex, Coup transcription factor α encoding, *Nr1d1* (t=7.259, *q*<0.001), and *Pgr* (t=5.275, *q*<0.001) were differentially expressed between female (n=10) and male (n=10) WT mice; **E)** Significance of sex-biased expression of nuclear receptor encoding genes in various human prenatal brain tissues. A1C: Primary auditory cortex; AMY: Amygdala; CBC: Cerebellar cortex; DFC: Dorsolateral prefrontal cortex; HIP: Hippocampus; ITC: Inferolateral temporal cortex; M1C: Primary motor cortex; MD: Mediodorsal nucleus of thalamus; MFC: Anterior cingulate cortex; OFC: Orbital frontal cortex; S1C: Primary somatosensory cortex; STC: Posterior superior temporal cortex; STR: Striatum; V1C: Primary visual cortex; VFC: V entrolateral prefrontal cortex. The dotted line marks nominal significance threshold (*p*<0.05). **F)** Illustration of BRD1 suggested role as a scaffold protein linking NRs with chromatin remodeling proteins, histone modifiers and histone acetyl transferases. Indicated BRD1 protein interaction partners are based on ^9,10^.

### Psychiatric disorder risk enrichment among genes containing NR binding motifs

Several NR binding neuro-active ligands (e.g. retinoic acid, vitamin D and thyroid hormone) have been associated with psychiatric disorders ^38–41^, but the contribution of their respective genomic actions in relation to psychiatric disorders risk is poorly understood. To address this matter, we used an *in silico* approach to assess enrichment for genetic disease risk in sets of genes containing putative NR promoter consensus sequences. Intriguingly, the predicted target genes of several NRs were significantly enriched with genetic schizophrenia risk (e.g. AR, HNF4G, VDR/RXRA, RARA/RXRG, RXRB, RAR-related orphan receptor (ROR) A, B, and C, RARA, and several types of ESRs) (**Figure 5C**). In addition, we found nominal significant enrichments for psychiatric disorder risk, including major depressive disorder in the target genes of RARA, RARA/RXRG, and LXRA/RXRA (**Figure 5C**). In a secondary analysis, we assessed whether NR target genes were enriched for common variant risk in diseases in which NRs are reportedly involved, namely type 2 diabetes (T2D) and rheumatoid arthritis. In line with peroxisome proliferator-activated receptor alpha (PPARA) ^42^, FXR ^43^ and ESR ^44^ agonists currently being used or have been suggested as treatment modalities of T2D, the target genes of PPARA/RXRA, FXR, ESR2 and ESRRA all showed nominal enrichment for T2D risk. Similarly in rheumatoid arthritis, vitamin A derivates have shown promising treatment effects and we found that RARA/RXRA and G and RXRB were nominally enriched with rheumatoid arthritis risk ^45^ **(Figure S9)**. Thus, these results supported the validity and relevance of the observed enrichments of psychiatric disorder risk.

### Sex-biased expression of nuclear receptor encoding genes in brain tissues

Sex-biased expression of the ASD candidate and NR encoding gene, *RORA*, has been suggested as a contributor to the sex-bias in ASD ^46^. We speculated that sex-biased expression of genes encoding NRs in general may be a contributing factor to the sex-biases in *Brd1*^+/−^ mice and in psychiatric disorders overall. Hence, we assessed expression of NR encoding genes in RNA-sequencing data on striatum and cortex from male and female WT mice. Out of 31 NR encoding genes expressed (RPKM > 1) in striatal tissue, expression of Coup transcription factors beta, 1, and 2 (*Nr1d2* (*q*<0.001), *Nr2f1* (*q*=0.002), and *Nr2f2* (*q*=0.026)), the progesterone receptor encoding gene, *Pgr* (*q*=0.032), and *Rxra* (*q*<0.001) differed significantly between female and male mice after Benjamini-Hochberg false discovery rate (FDR) correction at 5% (**Figure 5D**). Of the 32 NR encoding genes expressed in aCC, Coup transcription factor α encoding, *Nr1d1* (*q*<0.001), and *Pgr* (*q*<0.001) were differentially expressed between female and male WT mice **(Figure 5D**). To examine whether these sex-biased observations are corroborated in data from human brain, we assessed the overlap between NR encoding genes and reported sex-differentially expressed genes across brain regions at 4 developmental stages (prenatal, early childhood, puberty and adulthood) ^47^. We found that sex-biased genes are significantly enriched with genes encoding NRs at the earliest developmental stage, with particular enrichment in medial frontal cortex (*p*=0.006), orbitofrontal cortex (*p*=0.004) and in the striatum (*p*=0.01) (**Figure 5E**). The sex-differentially expressed NR encoding genes includes *NR1D1*, *NR2F2* and *RXRA*, as seen in adult mice, but additionally comprised genes encoding: thyroid hormone receptor beta (THRB); mineralocorticoid receptor (MR); NGF1-B, RORB; orphan receptors (NOR-1 and NURR1) among others at the prenatal stage (**Table S8**).

## DISCUSSION

Psychiatric disorders are heterogeneous and characterized by interconnected etiologies ^1,48^. Environmental and genetic risk factors are shared between diagnostic categories, and interact with each other in complex ways to influence phenotype ^49^. BRD1 has been implicated with both psychotic and affective disorders ^16–22^ and since it acts as an epigenetic regulator during neurodevelopment, it has the potential to integrate intrinsic and environmental signals into the shaping of the maturing brain. Here, we demonstrate that reduced *Brd1* expression in mice results in brain morphometric alterations accompanied by changes in behaviors and underlying neurobiology with broad translational relevance to psychiatric disorders. Interestingly, and mirroring the gender differences observed in psychiatric disorders ^2–6,50^, these changes are sex-biased with only female *Brd1*^+/−^ mice displaying changes in affective behaviors and only male mice showing increased sensitivity towards the psychotomimetic drug, cocaine. Adding to the growing understanding of BRD1’s molecular function, our study suggests that BRD1 acts as a modulator of NR mediated signaling and that dysregulation of subsets of NRs may significantly contribute to the pathological changes associated with reduced BRD1 expression. As NR mediated signaling is key to spatio-temporal transcriptional control during early neurodevelopment, we suggest that the sex-biased expression profile of these receptors may contribute to the sex-differential impact of reduced Brd1 expression in male and female mice and potentially to sex differences in mental disorders in general.

### Female, but not male, *Brd1*^+/−^ mice display behavioral changes with translational relevance to affective disorders

Cognitive impairments are common in psychiatric disorders, and although more thoroughly investigated in male *Brd1*^+/−^ mice ^26^, both sexes display cognitive impairments with broad translational relevance, including context-dependent learning deficits and impaired reference memory. PPI deficits, which has been linked to abnormalities of sensorimotor gating and have been reported in both schizophrenia and bipolar disorder ^51^, is similarly seen in both male and female *Brd1*^+/−^ mice along with exaggerated startle responsivity. However, neither male nor female *Brd1*^+/−^ mice display consistent changes in their risk-taking behaviors in the open field, bright open field or elevated plus maze. Female *Brd1*^+/−^ mice, additionally, did not exhibit marked changes in their circadian cycle and did not display increased sensitivity towards the psychomotor stimulatory effects of amphetamine and cocaine. However, supporting their translational value as model of depressive symptomatology seen in affective disorders ^52^ and the prodromal stage of schizophrenia ^53^, female *Brd1*^+/−^ mice displayed increased immobility during FST and TST indicating behavioral despair ^54^, and decreased sucrose preference representing anhedonia ^55^. Similar to what has been reported in both schizophrenia, bipolar disorder ^56^, and depressed suicide victims ^57^, female *Brd1*^+/−^ mice display abnormal neuronal morphology with reduced dendritic branching.

### Sex-biased neurochemistry and psychotomimetic drug sensitivity in *Brd1*^+/−^ mice

Sex differences in animal models of psychiatric disorders are common and may mirror the documented sex differences in psychiatric disorders where symptom profiles and severity differ between sexes ^2–6,50^ and where e.g. women are more susceptible to affective disorders than men ^2,5^. In line with the reported divergences in behaviors, the neurochemical profile of female *Brd1*^+/−^ mice varied significantly from what we have previously reported in male *Brd1*^+/−^ mice ^25,26^. Although both male and female mice had increased hippocampal dopamine, consistent with the monoamine hypothesis of depression ^58^, only female *Brd1*^+/−^ mice displayed significantly reduced levels of cortical serotonin and striatal dopamine. Correspondingly, male and female *Brd1*^+/−^ mice show markedly different regional expression of genes encoding neurotransmitter receptors, including receptors for dopamine and serotonin, which collectively may offer an explanation for the observed differences in sensitivity towards psychotomimetic drugs in female and male *Brd1*^+/−^ mice. Adding to this notion, female cortical DEGs were significantly enriched with major depressive disorder risk in line with their changes in affective behaviors, whereas male cortical and striatal DEGs, respectively, were enriched with, schizophrenia and bipolar disorder risk genes in accordance with their increased psychotomimetic drug sensitivity. Akin to suggested altered signal transduction cascades in both schizophrenia and affective disorders ^59–62^, calcium-, cAMP-mediated signaling and related signal transduction pathways were significantly enriched among cortical and hippocampal DEGs in female *Brd1*^+/−^ mice, and similar enrichments have been reported for DEGs in male mice ^25^. Despite their markedly different behavioral and neurochemical profiles, cortical DEGs in both sexes clustered in mitochondrial dysfunction and oxidative stress pathways, indicating common underlying effects of reduced *Brd1* expression in male and female mice. In line with this notion, variants in the mitochondrial genome have been associated with schizophrenia ^63^ and oxidative status has been linked with a range of psychiatric disorders ^64,65^.

### Global expression profiling suggests dysregulated nuclear receptor mediated signaling in *Brd1*^+/−^ mice

Interestingly, DEGs identified across brain regions in both male and female *Brd1*^+/−^ mice clustered in various pathways relating to NR mediated signaling. NRs and their ligands have been implicated with psychiatric disorders ^40,41,66–73^, including robust epidemiological association (e.g. between neonatal and maternal vitamin D status and schizophrenia ^74^ and autism spectrum disorder ^75,76^), and genome wide association of loci harboring the *RXRG* in bipolar disorder ^77^ and attention deficit hyperactivity disorder ^77^, *LXRA* in autism spectrum disorder ^78^, and *NR4A2* in major depressive disorder ^79^. Gene set and pathway analyses of GWAS data in major depressive disorder and bipolar disorder additionally reveal enrichment of genes implicated with thyroid hormone and retinoic acid signaling ^60^ as well as genes containing RAR/RXR consensus sequences ^79^. At the functional level, transcriptomic changes in post-mortem DLPFC samples from schizophrenia patients are enriched with NR encoding genes ^73^. NRs carry out their genomic functions through binding to co-activators and co-repressors which facilitate the recruitment of regulatory proteins, including histone deacetylase-, and acetyltransferase complexes ^80^. Interestingly, several co-regulators have been genome wide significantly associated with mental disorders (e.g. *BRD8, CNOT1, EP300, PAK6, KMT2E*, and *SLC30A9* in autism spectrum disorder ^81^ and/or schizophrenia ^78,82,83^*, KMT2D* and *GSN* in bipolar disorder ^84^ and/or schizophrenia ^85^, and *KDM4A* and *FOXP2* in ADHD ^86^). BRD1 is a multidomain protein and interacts with several transcriptional regulators identified in complexes with NRs ^87,88^, it’s interactome is significantly enriched with genes implicated with NR signaling ^10^, and BRD1 contains 5 putative NR binding sites ^34,35^. Other bromodomain containing proteins (TRIM24 ^89^, PB1 ^90^ and ATAD2 ^91^) have all been shown to directly interact with NRs and to mediate their ligand-dependent activation. As the RAR ligand, retinoic acid, is impaired in inducing differentiation in BRD1 depleted mouse embryonic stem cells ^15^, this collectively suggests that BRD1 facilitates a similar regulatory function on NR signaling (**Figure 5F**). Adding to this notion, multiple genes encoding key proteins implicated in neuro-active steroid bio-availability were dysregulated in the brains of female *Brd1*^+/−^ mice (see **Supplementary Discussion)**.

### Psychiatric disorders risk enrichment in the target genes of nuclear receptors and sex-biased expression of nuclear receptor encoding genes

NR mediated signaling has been associated with numerous pathologies such as cancer, inflammation, diabetes and lately psychiatric disorders ^40,41,66–73^ and NR mediated signaling has been suggested as a therapeutic target in schizophrenia ^72^. We show evidence suggesting that BRD1 modulates the genomic actions of NRs, and we provide *in silico* evidence that the target genes of several NRs are significantly enriched with psychiatric disorder risk. This is particularly the case for target genes of NR signaling pathways, which activity depend on BRD1 status in HEK cells and in mouse brain tissue (e.g. AR, HNF4, VDR, LXR/RXR, RAR, and estrogen receptor signaling), thus highlighting the translational relevance of *Brd1*^+/−^ mice as a model for psychiatric disorders. Brain development follows sex differential trajectories ^92^ with concordant regional sex-biased expression of comprehensive gene sets at various developmental stages. We show that this includes differential expression of genes encoding NRs particularly at the earliest stage of brain development in humans. Noticeably, the sex-differential regulation of NR encoding genes, is particularly seen in the same brain tissues (striatum and medial frontal cortex) that are associated with the behavioral and neurochemical changes that are sex-differentially affected in *Brd1*^+/−^ mice. As regional brain BRD1 expression does not appear to differ between men and women ^47^, it is thus likely, that hampered BRD1 availability affects the transcriptional control mediated by NRs at critical neurodevelopmental timepoints in a sex-specific manner. This in turn may result in differential changes in behavior and neurochemistry in the adult male and female *Brd1*^+/−^ mice.

Here we expand the behavioral, neuropathological and molecular characterization of a genetically modified mouse model that is based on the schizophrenia and bipolar disorder associated gene, *BRD1*. In line with a role for BRD1 as a scaffold protein linking histone modifiers and chromatin remodelers to transcriptional regulated sites at the genome, our data support a model in which BRD1 specifically modulate the genomic actions of NRs and their psychiatric risk enriched target genes. Combined with the accumulating epidemiological and genetic associations of nuclear receptors, their co-regulators and ligands to psychiatric disorders, our study adds to the growing evidence base supporting a central role for nuclear receptor mediated signaling in psychiatric disorders and their sex-bias.

## ACKNOWLEDGEMENTS

The study was supported by grants from The Danish Council for Independent Research– Medical Sciences (ADB and JHC), The Central Denmark Region (Region Midt) (JHC), The Augustinus Foundation (JHC), The Riisfort Foundation (JHC), The Lundbeck Foundation (ADB), The Faculty of Health Sciences, Aarhus University (APR, ADB), and The Novo Nordisk Foundation (ADB and JHC). Centre for Stochastic Geometry and Advanced Bioimaging was supported by Villum Foundation (JRN). We thank Tanja Stenshøj Østergaard for the genotyping of mice and Helene M. Andersen and Maj-Britt Lundorf for assisting in histology experiments.

## FINANCIAL DISCLOSURES

OM and ADB are coinventors on a patent application submitted by Aarhus University entitled “Method for diagnosis and treatment of a mental disease” (EP20060742417) that includes claims relating to *BRD1* among other genes. OM, ADB, JHC, MN, APR, and PQ are coinventors on a patent application submitted by Capnova A/S entitled “Genetically modified non-human mammal and uses thereof” (PCT/EP2013/069524) that includes the *Brd1*^+/−^ mouse. Besides being employed by H. Lundbeck A/S, KF, AM, and MD declares no biomedical financial interests or potential conflicts of interest. SHL, RL, JonP, NN, GuW, NL, VP, TF, JohP, BP, JRN, and GrW report no biomedical financial interests or potential conflicts of interest.

